# Animal septins contain functional transmembrane domains

**DOI:** 10.1101/2023.11.20.567915

**Authors:** Jenna A. Perry, Michael E. Werner, Shizue Omi, Bryan W. Heck, Paul S. Maddox, Manos Mavrakis, Amy S. Maddox

**Affiliations:** Department of Biology, The University of North Carolina at Chapel Hill; Chapel Hill, North Carolina, 27599 USA; Institut Fresnel, CNRS UMR7249, Aix Marseille Univ, Centrale Med, 13013 Marseille, France

## Abstract

Septins, a conserved family of filament-forming proteins, contribute to eukaryotic cell division, polarity, and membrane trafficking. Septins scaffold other proteins to cellular membranes, but it is not fully understood how septins associate with membranes. We identified and characterized an isoform of *Caenorhabditis elegans* septin UNC-61 that was predicted to contain a transmembrane domain (TMD). The TMD isoform is expressed in a subset of tissues where the known septins were known to act, and TMD function was required for tissue integrity of the egg-laying apparatus. We found predicted TMD-containing septins across much of opisthokont phylogeny and demonstrated that the TMD-containing sequence of a primate TMD-septin is sufficient for localization to cellular membranes. Together, our findings reveal a novel mechanism of septin-membrane association with profound implications for septin dynamics and regulation.

## Main Text

Cells comprise exceptionally complex mixtures of proteins, lipids, sugars, and small molecules that must combine to create multicomponent machines and to execute multistep signaling and synthesis cascades. The efficiency of such cellular tasks is often boosted by scaffolding via polymeric systems including the cytoskeleton. Septins are a highly conserved family of proteins that form palindromic hetero-oligomeric rods, which anneal into non-polar filaments and gauzes. Via their association with the plasma membrane, septin filaments recognize micron-scale membrane curvature, create diffusion barriers, and regulate cell morphogenic events via scaffolding other cytoskeletal polymers (i.e. F-actin and microtubules) and biochemical regulators of cell division, cell migration, and polarity establishment ^1,2^. While interaction with cellular membranes is thought to be crucial for septin polymer dynamics and function, how septins associate with membranes is not understood. Three polybasic regions (PB1, PB2, PB3) and an amphipathic helix (AH), are each sufficient for membrane interaction *in vitro*, and while the AH domain has been implicated in conferring membrane curvature sensing *in vivo* in the filamentous fungus *Ashbya*, the functionality of all of these domains in the context of intact septin complexes *in vivo* is still lacking ^3–9^. We explored how septins associate with membranes and discovered that some septins have functional transmembrane domains (TMDs).

### *C. elegans* UNC-61a contains a TMD that drives membrane localization

To study the mechanisms of septin-membrane association, we interrogated the uniquely simple set of septin genes of the nematode *Caenorhabditis elegans*. While other animal and fungal model organisms have five or more septin genes, often with many splice isoforms, the *C. elegans* genome contains only two septin genes, *unc-59* and *unc-61*. UNC-59 and UNC-61 proteins are usually interdependent *in vivo* and form palindromic tetramers *in vitro* ^10,11^. *C. elegans* septins exhibit behaviors shared by other organisms’ septins, including enrichment in and requirement for normal function of the cytokinetic ring ^10–13^. The *C. elegans* septin gene loci for *unc-61* encodes three isoforms: a, b, and c (Figure S1). *unc-61a* is transcribed from an alternative start site upstream of the shared start codon of isoforms *unc-61b* and *unc-61c*, which differ only by a single stretch of four amino acids, a distinction that we do not interrogate here. Hereafter, we refer to UNC-61b and UNC-61c collectively as UNC-61b/c. AlphaFold and a deep learning model for transmembrane topology prediction and classification (DeepTMHMM ^14^), predicted that the N-terminal amino acids of UNC-61a encode an α-helix that is expected with 99-100% certainty to form a transmembrane domain (TMD; Figure 1A and B). Alignment of homologous septin sequences from several related nematode species revealed that this putative TMD sequence is well conserved in this genus (Figure 1C). While the presence of a putative TMD has been predicted for at least 30 non-opisthokont (e.g. Chromista and Archaeplastida) septins ^15–17^, their relevance has not been tested experimentally. Importantly, to date, no TMD-containing septin (TMD-septin, hereafter) has been confirmed to be expressed within opisthokont (animal and fungal) lineages and functional implications of such a septin have not been investigated.

**Fig. 1.**
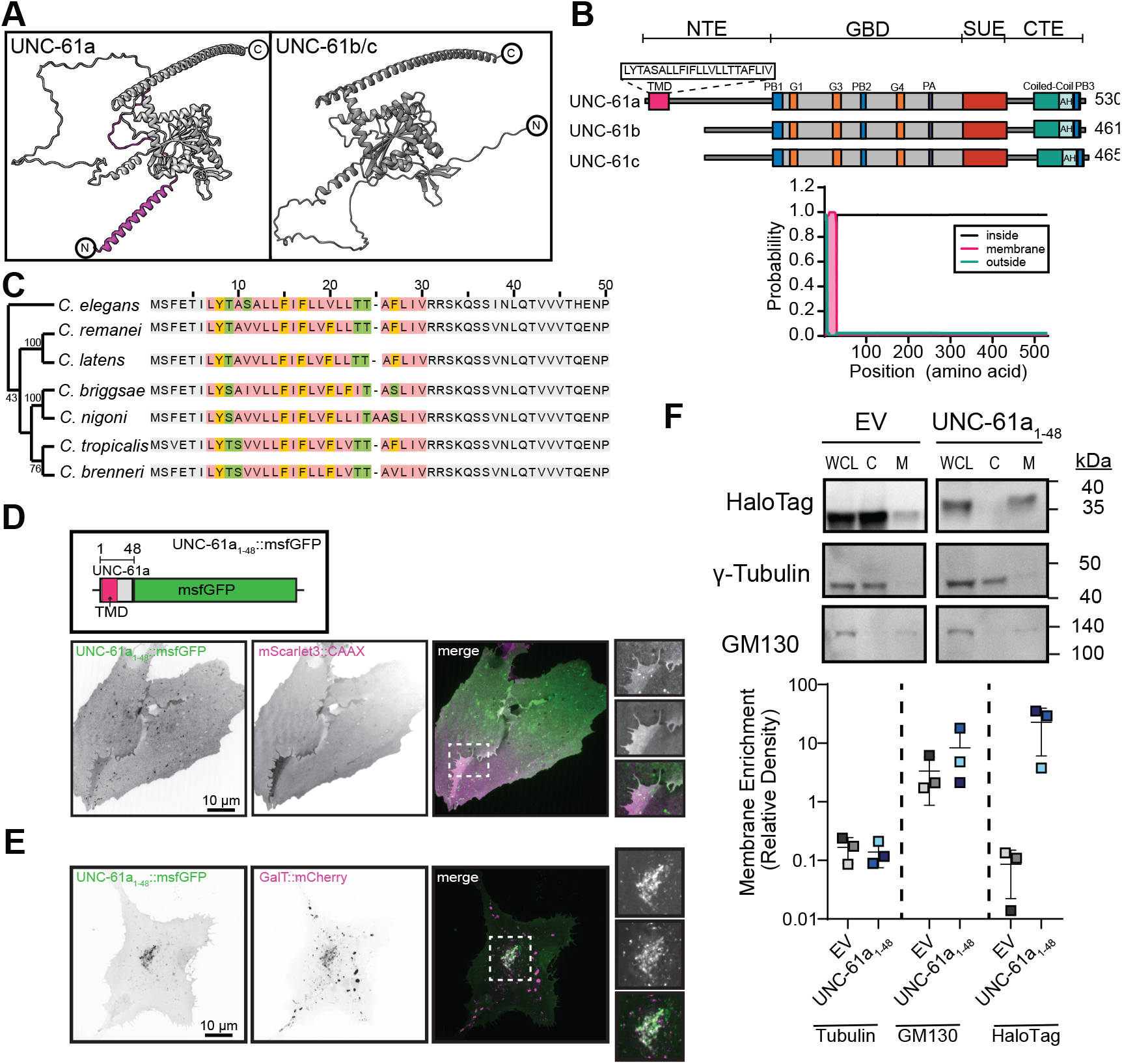
*C. elegans* UNC-61a contains a transmembrane domain. **(A)** AlphaFold structures of UNC-61a and UNC-61b/c. Magenta: N-terminal alpha-helix of UNC-61a. **(B)** (top) Illustration of UNC-61a. Motif colors: pink: transmembrane domain (TMD), grey: GTP-binding domain, red: SUE, blue: PB1/2/3, orange: GTPase domains, purple: polyacidic region (PA), teal: coiled-coil domain, light green: amphipathic helix (AH). Brackets at top of schematic denote septin domains (NTE: N-terminal extension, GBD: GTP-binding domain, SUE: septin unique element, CTE: C-terminal extension) (bottom) TMHMM output probability of TMD for *C. elegans* UNC-61. Teal: extracellular, pink: TMD, black: cytosolic. **(C)** (right) Alignment of the first 50 amino acids of UNC-61 homologs in *C. elegans* and closely related nematodes. Non-TMD residues: grey, TMD sequence: colored based on amino acid properties (hydrophobic: rose, aromatic: orange, positively charged: blue, negatively charged: red, hydrophilic: green, cystine: yellow). (left) Neighbor-joining tree of evolutionary relationships. **(D)** (top) Illustration of UNC-61a _1-48_::msfGFP construct, highlighting the position of msfGFP (green), TMD (magenta), and UNC-61a fragment (grey). (bottom) U2OS cells transiently co-expressing UNC-61a TMD_1-48_::msfGFP (green: and plasma membrane label mScarlet3::CAAX( magenta). Images shown are a single plane, inverted contrast, and scaled to display minimum and maximum pixel intensities. Dashed box (white) indicates region used for 2x inset. Scale bar = 10 μm. (E) U2OS cells transiently co-expressing UNC-61a TMD_1-48_::msfGFP (green) and GalT::mCherry (Trans-Golgi and Golgi transport intermediates; magenta). Images are maximum intensity projections, inverted contrast, and are scaled to display pixel intensity ranges. Dashed box (white) indicates region used for 2x inset. Scale bar = 10 μm. **(F)** (top) Western blots of cytosolic (C) and membrane (M) fractions of HeLa cells transiently expressing either HaloTag empty vector (EV) or UNC-61a TMD_1-48_::HaloTag (UNC-61a TMD_1-48_). Lanes 1-3: whole cell lysate (WCL) blotted singly with antibodies recognizing GM130 (membrane), γ-tubulin (cytosol) and HaloTag. Lanes 4-6 WCL,C, and M fractions are probed with a mixture of all three antibodies (GM130, γ-tubulin, and HaloTag). A shorter exposure (4.9 s) was used to image the HaloTag to prevent band saturation. (bottom) Membrane enrichment of HaloTag for the EV and UNC-61a TMD_1-48_::HaloTag was analyzed semi-quantitatively and plotted as the ratio of membrane enrichment to cytosolic enrichment. Plot: mean ± SD of three independent experiments.

To test the functionality of the putative TMD of *unc-61a*, we assessed whether this sequence was sufficient to drive membrane association. We created a transgene encoding residues 1-48 of UNC-61a (the unique first exon of *unc-61a*) and msfGFP (Figure 1D) and co-transfected U2OS cells with UNC-61a_1-48_::msfGFP and either mScarlet3::CAAX to label the plasma membrane or GalT::mCherry, which labels the trans-Golgi and Golgi transport intermediates, (Figure 1D,E). UNC-61a_1-48_::msfGFP localized to both the plasma membrane and the trans-Golgi indicating that the putative TMD does localize with membranes and membrane-bound compartments (Figure 1D,E). To further test if the UNC-61a_1-48_ peptide directly interacted with cellular membranes, we created a transgene with UNC-61a_1-48_ fused to a HaloTag, transfected the construct into HeLa cells and separated HeLa cells into cytoplasmic and membranous fractions (Figure 1E, S2). In contrast to γ-tubulin, which is enriched in the cytoplasmic fraction, UNC-61a_1-48_::HaloTag was enriched in the membrane fraction and was undetectable in the cytoplasmic fraction, as was the Golgi matrix protein GM130. These findings demonstrate that the predicted TMD of *C. elegans* UNC-61a can confer localization to membranes.

### UNC-61a TMD is responsible for UNC-61a function in vulval morphology maintenance

To test whether a septin bearing a TMD is functionally important, we first examined how the localization of UNC-61a compared to that of UNC-61b/c and UNC-59. To minimize interference with the N-terminal TMD, we generated animals expressing UNC-61a intramolecularly tagged with a *C. elegans* codon optimized GFP and placed under the control of the *unc-61a* promoter at a Mos1-mediated single copy insertion (MosSCI) site on chromosome II (Figure S3). A similar MosSCI approach was utilized to create an exogenous copy of UNC-61b/c tagged at its N-terminus with the monomeric and *C. elegans* codon optimized wrmScarlet under control of the *unc-61b/c* promoter at a MosSCI site also on chromosome II (wrmScarlet::*unc-61b/c;* Figure S3). UNC-61a::GFP was highly enriched in the terminal bulb of the pharynx, often decorating linear structures, and in the vulva in adult hermaphrodite *C. elegans*, while wrmScarlet::UNC-61b/c was less enriched in those structures (Figure 2A). Conversely, wrmScarlet::UNC-61b/c decorated the *C. elegans* oogenic germline; UNC-61a::GFP was undetectable there (Figure 2A; ^11,18^). Interestingly, fluorescently-tagged UNC-59 (UNC-59::GFP) expressed from its endogenous locus localizes to tissues where either UNC-61a or UNC-61b/c is individually enriched: UNC-59::GFP is present in pharynx, germline, and vulva (Figure 2A). The differential localization patterns of the UNC-61 isoforms suggest that they have distinct physiological roles in the tissues where they are expressed and that UNC-59 functions with both UNC-61a and UNC-61b/c, in different tissues.

**Fig. 2.**
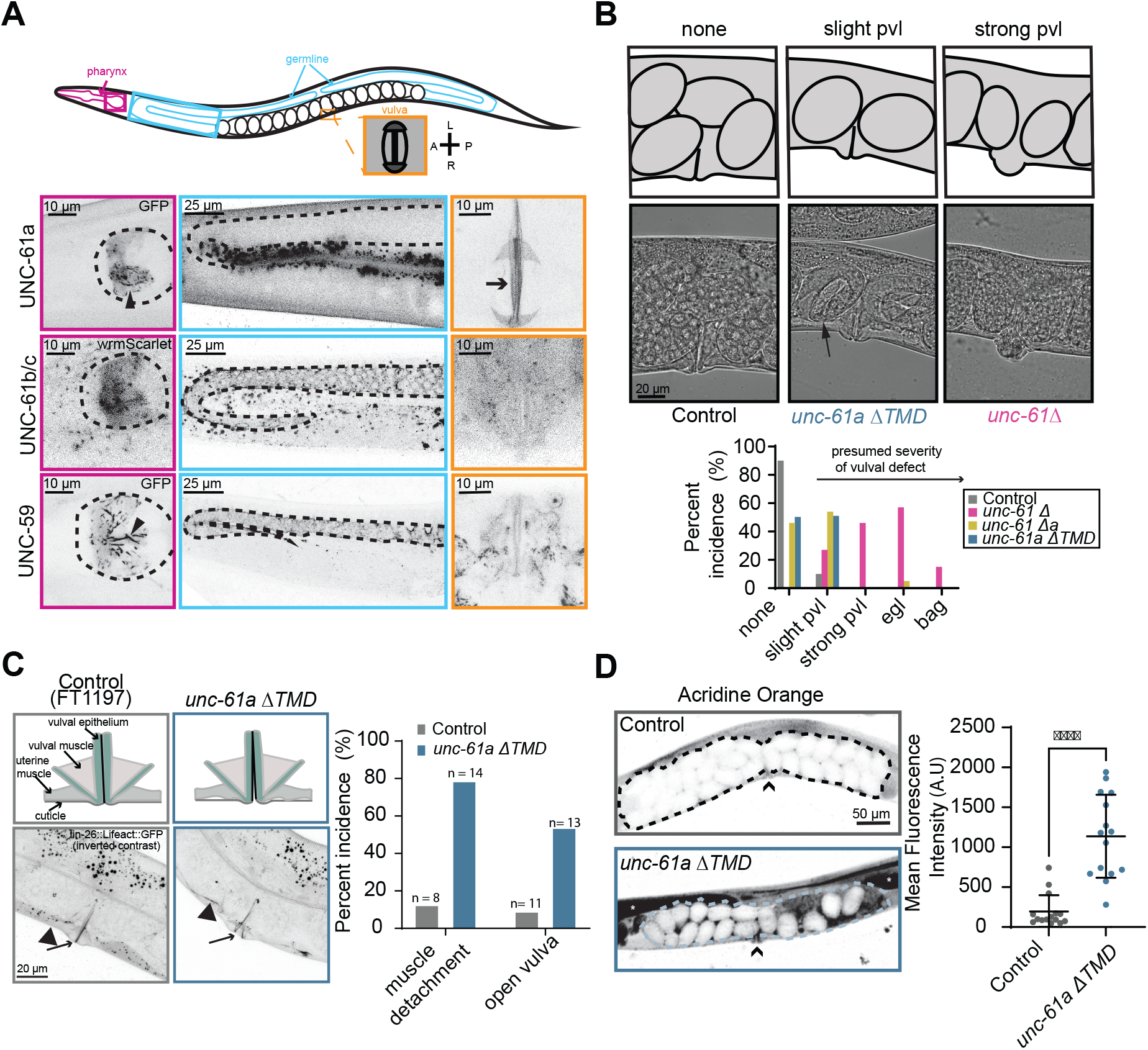
UNC-61a TMD is required for normal vulval morphology and function. **(A)** (top) Schematic of adult *C. elegans* hermaphrodite (pink: pharynx, blue: germline, orange: vulva.) (bottom) Images are inverted contrast of the fluorophore annotated. Arrow: vulval epithelium apical surfaces; arrowhead; linear septin decorated structures. Dashed lines outline the *C. elegans* terminal bulb and oogenic germline. The fluorescent signal beneath the germline corresponds to the autofluorescence of the *C. elegans* intestine. Scale bar = 10 μm (pharynx and vulva); 25 μm (germline). **(B)** (top) Schematic of representative images of protruded vulva (pvl) phenotypes and embryos in uterus. (middle) Representative transmitted light images of pvl phenotypes. Arrow denotes embryos that are more developed than they should be indicating an egg-laying defective (egl) phenotype. (bottom) Percent incidence of post-embryonic phenotypes (pvl: protruded vulva; egl: egg-laying defective; bag: bag-of-worms) of control (grey), *unc-61Δ* (pink), *unc-61 Δa* (yellow), and *unc-61a* ΔTMD (blue) animals (n=100 animals). **(C) (**left top) Schematic of control and *unc-61a* Δ*TMD* vulvae (green: epithelium, black: apical surface, grey: surrounding muscles.) (left bottom) Inverted contrast images of Lifeact::GFP (F-actin) in vulva region. Arrowhead: muscle cuticle connection in control, disconnection in *unc-61a* ΔTMD; arrows: apical surface of the vulval epithelium. (right) Percent incidence of muscle detachment and open vulva phenotypes. “n” above bar indicates the number of animals scored for each condition. Scale bar = 20 μm. **(D)** (left) Representative images of uterine and embryo permeability in control and *unc*-*61a ΔTMD* animals. Arrowheads denote vulval location, asterisks indicate germline location, and dashed lines outline uterus. Scale bar = 50 μm. (right) Quantification of uterine (dashed lines) mean fluorescence intensity in control (grey; n = 15 animals) and *unc*-*61a ΔTMD* animals (blue; n = 15 animals). Error bars are mean ± SD. ****, ≤ 0.0001, as determined by Student’s unpaired t-test.

We next tested whether the loss of the TMD from UNC-61a affected its function in *C. elegans*. Depletion or mutation of either *unc-59* or *unc-61* results in uncoordinated movement, reduced fertility, and gross morphological defects of the egg-laying apparatus (uterus and vulva) ^11,18–22^. UNC-61 depletion and mutations used to date altered or removed expression and function of all three isoforms ^11,23–25^. We used CRISPR/Cas9 genome editing at the endogenous locus to excise amino acid residues 1-524 (numbering with respect to *unc-61a* translation) to create *unc-61* null animals, introduced a stop codon at amino acid residue 2 of *unc-61a* to create *unc-61a* null animals (*unc*-61Δa; Figure S4), and excised the bases encoding amino acids 7-29 to create a genotype specifically lacking the TMD of UNC-61a (*unc-61a* ΔTMD; Figure S4). *unc-61* null (*unc-61*Δ) animals exhibited the range and severity of phenotypes, including the retention of embryos in the uterus (egl) and larvae present inside the uterus (bag) (Figure 2B), that were previously reported for random mutagenesis-generated hypomorphic alleles ^11,18,19,22,23,26^. While *unc-61a* null (*unc*-61 Δa) or *unc-61*a ΔTMD animals had normal fertility and animal motility (Figure S5), they exhibited abnormally protruded vulvae (pvl; Figure 2B). Thus, the array of phenotypes exhibited by animals lacking all UNC-61 isoforms is partially due to absence of UNC-61b/c (fertility and motility defects) and partially due to the absence of UNC-61a (vulval morphology), whose function depends on its TMD, as excision of the TMD phenocopies the *unc-61a* null allele.

To understand the basis for the pvl phenotype in *UNC-61*a ΔTMD animals, we examined vulval morphology. High resolution imaging of adult animals expressing Lifeact::GFP to label F-actin in epidermal cells revealed that the vulva of the UNC-61a ΔTMD animals rested in an abnormal, open conformation (Figure 2C). In addition, the surrounding tissue frequently appeared detached from the ventral cuticle (Figure 2C). To test the possibility that these abnormalities of vulval morphology correspond to decreased barrier integrity, we exposed control and UNC-61a ΔTMD animals to acridine orange, a fluorescent dye whose internalization is blocked by the cuticle ^27,28^. In control animals, acridine orange did not enter the uterus or intrauterine embryos, while the uterus and some embryos in UNC-61a ΔTMD animals were strongly labeled, indicating vulval and extracellular matrix permeability (Figure 2D). Together with the prominent enrichment of UNC-61a on the vulval lumen (Figure 2A), these results suggest that UNC-61a via its TMD contributes to vulval morphology and therefore animal impermeability. In sum, our work demonstrates the functional importance of a TMD-septin.

### Septins throughout phylogeny are predicted to contain transmembrane domains (TMD)

Given the conservation of sequence and function of septins, we reasoned that TMD-septins could be present throughout opisthokont phylogeny. We surveyed over 34,000 sequences containing a “septin-type guanine nucleotide-binding (G) domain” on UniProt for the presence of TMDs ^29^. Putative TMDs were predicted to be present in septin genes from many organisms across phylogeny (Figure 3A). Of the 2320 eukaryotic species predicted to express septin-like proteins, 447, or nearly 20%, were predicted to possess at least one TMD-septin. TMD-septins were not predicted in many well-established model organisms including budding yeast, fruit fly, and mouse, but were predicted in some organisms, including the small primate *Carlito syrichta*, a non-model-animal tarsier (Figure 3A).

**Fig. 3.**
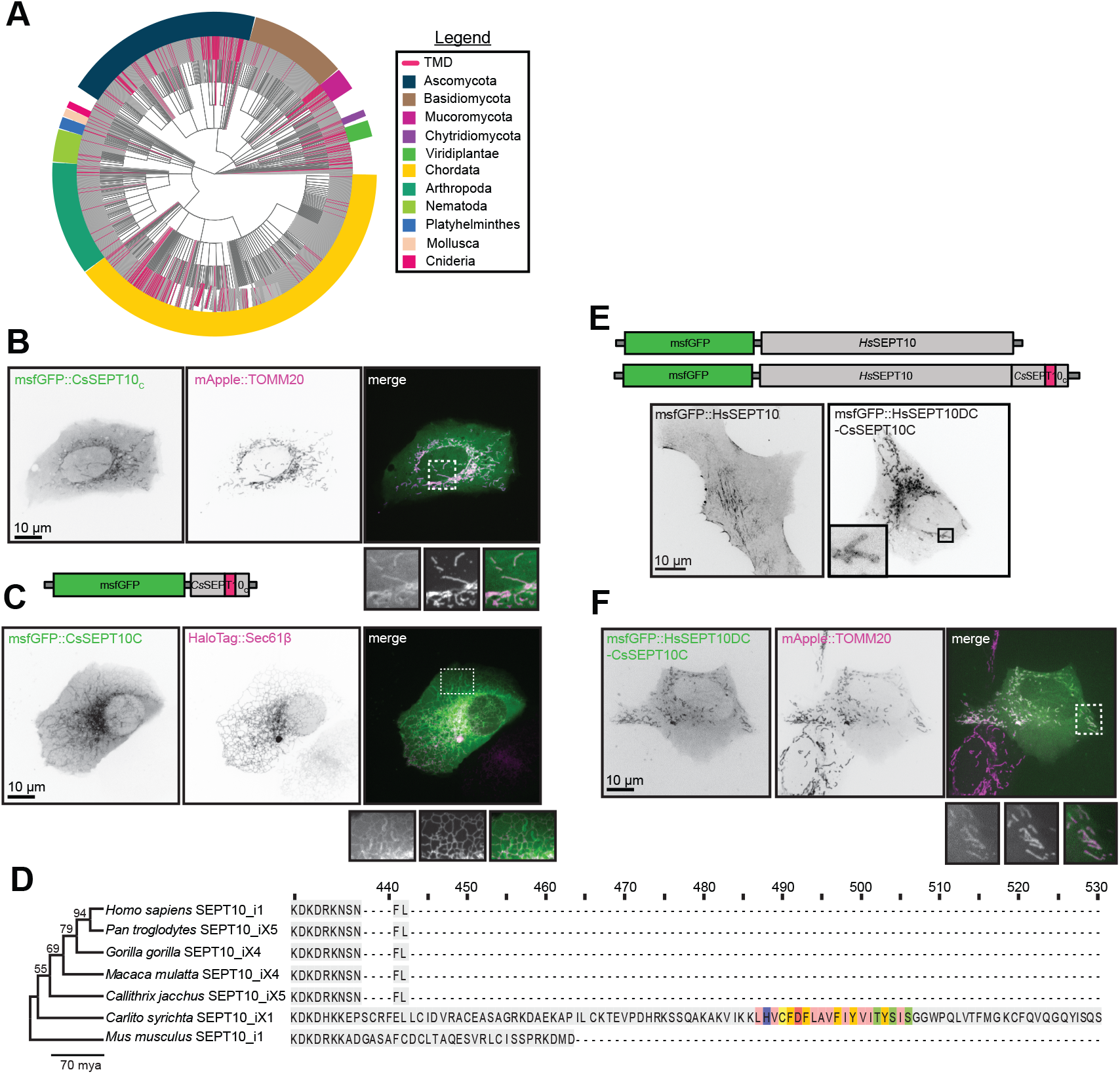
Animal septins are predicted to contain transmembrane domains (TMD). (A) Cladogram of all septin sequences deposited on UniProt (magenta: sequences containing TMDs). Rim colors: organismal phyla (dark blue: Ascomycota, brown: Basidiomycota, fuchsia: Mucoromycota, purple: Chytridiomycota, lime green: Viridiplantae, yellow: Chordata, teal: Arthropoda, light green: Nematoda, blue: Platyhelminthes, light pink: Mollusca, hot pink: Cnideria **(B)** msfGFP::*Cs*SEPT10_C_ (green) expressed in U2OS cells and colocalized with Translocase of Outer Mitochondrial Membrane 20 (TOMM20; magenta). **(C)** msfGFP::*Cs*SEPT10_C_ (green) co-expressed with HaloTag::Sec61β (ER; magenta) in U2OS cells. **(D)** (Left) Neighbor-joining tree of evolutionary relationships (Right) Schematic of *Cs*SEPT10 and *Hs*SEPT10 protein architecture. Colors as in Figure 1C. **(E)** Localization of msfGFP::*Hs*SEPT10 and the chimera msfGFP::*Hs*SEPT10_ΔC_-*Cs*SEPT10_C_. **(F)** Colocalization of the chimera msfGFP::*Hs*SEPT10_ΔC_-*Cs*SEPT10_C_ (green) and mApple::TOMM20 (magenta). All images are inverted contrast, unless merged, and insets are magnified 2x. Scale bar = 10 μm.

To test whether the predicted TMD of a primate septin is sufficient for localization to membranes, we examined *C. syrichta* SEPT10 (*Cs*SEPT10) whose C-terminus contains a TMD sequence (Figure S6). We created a fusion protein wherein a fluorescent protein was added to the TMD-containing C-terminus of the *Cs*SEPT10 and expressed GFP::*Cs*SEPT10_C_ in U2OS cells (Figure 3B). The predicted TMD was sufficient to localize the fluorescent protein to the outer mitochondrial membrane, as indicated by the colocalization of CsSEPT10 and the mitochondrial translocase TOMM20, and to the endoplasmic reticulum membrane, as indicated by the colocalization of *Cs*SEPT10 with Sec61β. (Figure 3B and 3C). We next tested whether the TMD of *Cs*SEPT10 could drive membrane localization of a septin lacking TMD. Despite the close phylogenetic relationship between humans and *C. syrichta, Hs*SEPT10 is not predicted to contain a C-terminal TMD (Figure 3D, Figure S5). Accordingly, *Hs*SEPT10 localizes to actomyosin stress fibers (Figure 3E; ^30,31^). We created a fluorescently labeled chimeric SEPT10 by adding the *C. syrichta* C-terminal TMD to *Hs*SEPT10 (Figure 3F). The *C. syrichta* SEPT10 TMD was sufficient to re-localize *Hs*SEPT10 entirely from stress fibers to mitochondria (Figure 3E and F). Together, these results demonstrate that primates possess a TMD-septin. Furthermore, the presence of the TMD in septin genes throughout phylogeny suggests that an ancestral septin had a transmembrane domain that has been lost in some lineages and over the course of gene duplications.

## Discussion

Here, we report the first functional characterization of a TMD-containing septin, using the *C. elegans* UNC-61 isoform, UNC-61a, that contains a TMD that is highly conserved among nematodes. UNC-61a is expressed in vulval epithelial cells and in a subset of cells in the terminal bulb of the pharynx. Loss of UNC-61a resulted in defective vulval morphology, which can be attributed to the loss of the TMD. The UNC-61a TMD is sufficient to localize a fluorophore to cellular membranes. Additionally, the TMD from an isoform of SEPT10 from the primate *C. syrichta* localized a fluorescent probe and the actin-associated human SEPT10 to cellular membranes. Our work implicates the TMD in targeting septins to membranes, a novel mechanism for septin-membrane interaction.

To explore the evolutionary conservation of TMD-containing septins, we searched among 34,000 predicted septin genes for predicted TMDs and identified roughly 700 sequences from 450 organisms across opisthokont phylogeny. While the full list of predicted TMD-septins awaits experimental validation, our findings indicate a sporadic distribution of these proteins within phyla. Several factors could contribute to this pattern: current TMD prediction algorithms may fail to identify some TMD sequences (e.g. short helices do not conform to current criteria). Additional TMD-containing septins could be missed or misidentified due to the incompleteness of genome or transcriptome assemblies. Alternatively, the intermittent appearance of TMD-septins could reflect sporadic loss during the evolution from an ancestral septin that possessed the TMD. To understand the evolutionary history of TMD-septins, further advancements in genome annotation and transmembrane domain prediction tools will be crucial.

The TMD of *C. elegans* and related nematodes’ UNC-61a is N-terminal to the GTP-binding domain, yet other organisms’ septins contain a C-terminal TMD. Given the evolutionary distance among species encoding a TMD-septin, this inconsistency is not unprecedented and may indicate exon shuffling, wherein exons that correspond to functional gene domains are duplicated, changed, and rearranged to give rise to proteins with new functions while maintaining established roles ^32–34^. Further analysis of the DNA sequence surrounding the TMD will help test whether exon shuffling occurred in the evolution of TMD-septins. Additionally, previous work exploring the variability of the C-terminal extensions (CTE) in mitotic yeast septins uncovered functional differences related to the recruitment of non-septin proteins ^35^. It is possible that TMD septins coupled to a membrane via the N terminus versus the C terminus have distinct cohorts of interacting protein based on what regions of the protein are occluded or exposed.

To date, two motifs (the amphipathic helix (AH) and polybasic domains 1, 2, and 3 (PB1/2/3)) have been identified to facilitate septin-membrane interactions. AHs sense membrane curvature and PB1/2/3 confer lipid binding specificity ^7,36,37^. Disruption of these domains, however, is not sufficient to perturb septin localization to cellular membranes ^3–7,36^. This may be attributed to septins’ indirect association with the plasma membrane via other cytoskeletal components, such as the scaffold protein anillin that directly binds the plasma membrane ^26,38–40^. The presence of a validated TMD sufficient for membrane interaction in a multicellular organism prompts the reconsideration of mechanisms through which septins interact with cellular membranes. Furthermore, membrane association via a TMD is predicted to be more stable than association via an AH or via PB1/2/3. The majority of septin biology has been studied using species that appear not to express TMD-septins. Future work will examine whether the septins of a given species or cell type use one or more of the now three mechanisms of membrane association.

The *C. elegans* TMD-septin UNC-61a localizes to distinct regions of the adult animal: the terminal bulb of the pharynx and the apical membrane of vulval epithelial cells. Interestingly, a key function of these highly dynamic tissues is to secrete extracellular matrix ^41,42^. Our data assessing cuticle permeability indicate that loss of the TMD from UNC-61a reduces the integrity of vulval barrier function. The vulva may remain in an open configuration in UNC-61a ΔTMD animals due to a defect in secretion at the vulval epithelial cells’ apical surface ^43–48^. The extracellular matrix component O-glycan is highly enriched on the vulval lumen and under the flaps of cuticle that flank the vulval slit, in two half-circles where UNC-61a is uniquely enriched ^49^. Future work will explore the relationship between TMD-septins and extracellular matrix deposition.

Our characterization of UNC-61a prompts a reconsideration of the fundamental unit of septin polymerization in *C. elegans*. We propose that the septin heteromer is not always a tetramer of two copies of UNC-59 and two copies of UNC-61b/c. Rather, UNC-61a may replace one or both copies of UNC-61b/c in some cell types. Variability in the stoichometry of septin oligomers has precedent in human septin heteromers, which can be hexamers or octamers, and can co-polymerize into heterogeneous filaments ^50^. Similarly, tubulin isotypes with low expression can alter microtubule polymer dynamics, with cellular and tissue-level implications, such as in cancer prognosis ^42,51– 54^. We predict that TMD-septins, even when present as minor components of a septin system, interspersed into septin filaments, can have profound effects on septin functions, such as membrane association. Reconstitution of septin cohorts including TMD-septins and genetic tuning of septin expression in *C. elegans* will help elucidate biochemical and functional characteristics of TMD-containing filaments.

## Methods

### Phylogenetics

To find transmembrane domain (TMD) containing septins, we searched UniProt ^29^ using accession code IPR030379 to identify putative septins based on the presence of a Septin-type guanine nucleotide-binding (G) domain. Then, the resultant list was filtered for the “Transmembrane” annotation in UniProt. This filter uses the SAM (Sequence Analysis Methods) for automatic annotation to predict putative transmembrane domain when there is an overlap of at least 10 amino acids between TMHMM and Phobius predictors ^55,56^. The filtered list hits were parsed for duplicates, then organisms containing TMD-septins were plotted on a phylogenetic tree using Interactive Tree of Life (iTOL) and customized with R package ggTree ^57^. . Protein sequence alignments were conducted using MUSCLE ^58^ in MEGA11 ^59^. A neighbor-joining tree was generated with 1000 bootstrap replicates for statistical support in MEGA11.

### C. elegans strains and culture

*C. elegans* strains were maintained at 20 °C using standard culturing conditions ^19^. The strains used and/or generated for this work are listed Table 1.

### Human cell culture

HeLa cells were maintained in DMEM with 10% Fetalplex and 1% Pen/Strep. U2OS cells were maintained in McCoy’s medium (Thermo Fisher, Cat #16600082) supplemented with 10% fetal bovine serum (Dominique Dutscher, Cat # S181H), and 100 U/mL penicillin and 100 μg/mL streptomycin (Sigma-Aldrich Cat#P4333). Both cell types were cultured at 37 °C in 5% CO_2_.

### Mammalian plasmid construction and transient cell lines

UNC-61a TMD_1-48_:HaloTag was cloned into a backbone containing a CAG promoter, IRES, and hygromycin resistance cassette. HeLa cells were transfected with either the Double UP Halotag to mScarlet (a gift from Erik Dent; Addgene plasmid # 125138) or UNC-61a TMD_1-48_:HaloTag using Effectene (Cat# 301425, QIAGEN) according to the manufacturer’s instructions. Cells were seeded at 8.0 × 10^5^ the day prior to transfection in a 10 cm tissue culture dish. A total of 2 μg plasmid DNA and a 1:10 ratio of DNA to Effectene Reagent were used per transient transfection.

*Hs*SEPT10_i1 cDNA (UniProt Q9P0V9) was a gift from Andrew Wilde (University of Toronto). Synthetic msfGFP::*Cs*SEPT10_C_ and *Hs*SEPT10_i1_DC_-*Cs*SEPT10_C_ were from Eurofins Genomics (Germany). mScarlet3 DNA was from Addgene (plasmid #189753). CMV-driven msfGFP::*Hs*SEPT10_i1 (pM321), msfGFP::*Cs*SEPT10_C_ (pM324) and msfGFP::*Hs*SEPT10_i1_DC_-*Cs*SEPT10_C_ (pM325) and mScarlet3-CAAX (pM300) were cloned with seamless cloning (Cat# 638910, In-Fusion HD Cloning Plus Kit from Takara Bio) using NheI/BamHI linearized pCMV backbones (Clontech) and the oligonucleotide primer sequences are listed in Table 2.

The C-terminal sequence of *Cs*SEPT10 (*Cs*SEPT10_C_) is fused to the C-terminus of msfGFP with a 12-residue linker (GGGGSGGGGSSG) and comprises the 106 C-terminal residues of *Cs*SEPT10 i.e., it starts at NLRKDKD. The chimera *Hs*SEPT10_i1_DC_-*Cs*SEPT10_C_ contains this very same sequence of *Cs*SEPT10 i.e., *Hs*SEPT10_i1_DC_ misses its seven terminal residues, and the sequence of the fused region is NLRKDKDHKKE (Figure S5).

UNC-61a TMD_1-48_::msfGFP was cloned with seamless cloning ( NEBuilder HiFi DNA Assembly Master Mix, Cat# E2621) using NheI/BamHI linearized and calf intestinal alkaline phosphatase (NEB, Cat# M0525)-treated pM324 backbone. Fragments bearing 20bp overhangs with homology to the flanking regions were generated via PCR with Q5 High-Fidelity DNA Polymerase (NEB, Cat #M0491S) and gel purified (Monarch® Spin DNA Gel Extraction Kit, NEB Cat# T1120) prior to assembly. The oligonucleotide primer sequences are listed in Table 2.

U2OS cells were either single-transfected (pM321) or co-transfected (pM324pM325 or, pJAP11) with organelle markers using jetPRIME (Cat# 101000015, PolyPlus) according to the manufacturer’s instructions. TOMM20::mApple (outer mitochondria membrane) was from Addgene (plasmid #54955), GalT::mCherry (Golgi membrane) was from Addgene (plasmid #55052) and HaloTag::Sec61β (ER membrane) was a kind gift of Christopher Obara (UCSD). Cells were seeded at 3.0 × 10^4^ the day prior to transfection in a 24-well glass bottom plate (Cellvis, P24-1.5H-N). 0.2 μg of each plasmid DNA and a 1:2 ratio of DNA to jetPRIME Reagent were used per transient transfection. 24 hours post transfection, the culture medium was exchanged by Leibovitz medium (Cat# 21083027 Gibco) supplemented with 10% fetal bovine serum and antibiotics. Cells transfected with HaloTag::Sec61β were incubated in Leibovitz medium containing 250 nM JF585-HaloTag ligand (kind gift of Luke Lavis, Janelia) for 30 min and washed twice for 5 min with dye-free medium.

### Fluorescence imaging conditions and analysis

#### C. elegans

*C. elegans* fluorescence imaging was performed as described ^22^. Whole worm images were acquired using a Nikon A1R microscope with a 60 × 1.27 NA Nikon water immersion objective with a gallium arsenide phosphide photo-multiplier tube (GaAsP PMT) using NIS-Elements software. 25-30 confocal sections were acquired with 1 μm z-spacing.

#### U2OS cells

Cells were imaged live using a spinning disk unit (CSU-X1-M1 from Yokogawa) connected to the side-port of an inverted microscope (Eclipse Ti2-E from Nikon Instruments) using a Nikon Plan Apo ×100/1.45 NA oil immersion objective lens, a 488 nm and 561 nm laser line (Coherent) and an iXon Ultra 888 EMCCD camera (1024×1024 pixels, 13×13 μm pixel size, Andor, Oxford Instruments). Cells were maintained at 37 °C in a heating chamber (OkoLab H301-T-UNIT-BL).

### Cell fractionation assay and western blotting

Cell fractionation was performed using Mem-PER™ Plus Membrane Protein Extraction Kit (ThermoFisher Scientific, Cat # 89842) according to manufacturer instructions. In brief, 10 × 10^6^ cells were harvested and washed 3 times with Cell Wash Solution, after which 5 × 10^6^ cells were saved for whole cell lysate. The remaining 5 × 10^6^ cells were resuspended in Permeabilization Buffer and incubated at 4 °C with constant mixing. Permeabilized cells were centrifuged at 16000 × *g* for 15 minutes and the supernatant containing the cytosolic fraction was removed and stored in Laemmli buffer at -20 °C. The cell pellet was resuspended in Solubilization Buffer and incubated at 4 °C with constant mixing. Following incubation, the solubilized cells were centrifuged at 16000 × *g* for 30 minutes and the supernatant containing solubilized membrane and membrane-associated proteins was removed and stored in Laemmli buffer at -20 °C.

Fractions were resuspended in Laemmli buffer for a concentration of 6.7 × 10^3^ cells per μl and boiled for 10 minutes. The lysate equivalent of 8 × 10^4^ cells per lane was separated using anyKD Mini-PROTEAN TGX precast protein gels (Bio-Rad Laboratories, CA) and transferred to PVDF membrane (BioRad, Cat # 1620177). Membranes were blocked for 1 hr at RT with 5% (w/v) non-fat dried milk dissolved in TBS containing 0.1% Tween-20 (TBST). Membranes were incubated with primary antibodies diluted in blocking buffer for 1 hr at RT. The membranes were then washed three times in TBST and incubated for 1 hr at RT with secondary antibodies conjugated to horseradish peroxidase (GE Healthcare, Cat #NA931V). Following three washes in TBST, membranes were imaged using Amersham ECL Select Western Blotting Detection Reagent (GE Healthcare, Cat # RPN2235) and a BioRad Gel Doc XR+ system.

### C. elegans MosSCI transgenesis

Transgenes were injected into *C. elegans* strain COP93, along with constructs encoding for the Mos transposase and four negative selection markers against extrachromosomal plasmid arrays. The promoter sequence used for UNC-61a::GFP includes 690 bp of upstream sequence including the last exon of the upstream gene Y50E8A5.1. UNC-61a::GFP was internally tagged between the first and second exon of UNC-61a, which is a region unique to this isoform. The promoter sequence of wrmScarlet::UNC-61b/c comprised of the 524 bp upstream of its start codon. While this region does include part of the transcript for UNC-61a, the start codon was mutated to isoleucine to prevent read-through. A sequence encoding (Gly)_3_Ser(Gly)_4_Ser linker was added 3’ to the fluorescent protein sequence in both constructs. Integrants were selected for rescue of uncoordinated phenotype (Unc) of the parental strain and confirmed by PCR using the primers found in Table 3.

### C. elegans CRISPR/Cas9 genome editing and sequencing

CRISPR/Cas9 was performed as described ^60^. Briefly, a mix containing crRNA, tracrRNA, and Cas9 protein with a *dpy-10* co-CRISPR was injected into the gonad of young adult animals. Benchling electronic lab notebook ^61^ was used to select the optimal Cas9 cleavage site, the optimal sequence for the crRNAs, and to predict off-target possibilities (Table 4). The edited locus was amplified via PCR from *C. elegans* genomic DNA extracted from 8-10 adult worms using the Extract-N-Amp^TM^ Tissue PCR Kit (Millipore Sigma, MA) as previously described ^62^ and sequenced (Table 4). The resultant worm strains were outcrossed at least three times to N2 wildtype animals.

### Brood size

10 fourth larval stage (L4) animals were selected from a non-starved growth plate and placed on individual culture plates at 20 °C for 48 hours, after which the adult animal was removed. After two days, the plates with larval offspring were moved to 4 °C for 20 minutes to halt worm movement. All larvae and unhatched embryos were manually counted using a clicker on a Nikon SMZ800 dissection microscope.

### Plate level phenotype analysis

To score plate-level phenotypes, 100 adult animals from the brood size measurement culture plates (described above) were manually assessed and scored using a Nikon SMZ800 dissection microscope.

### Thrashing assay

Animal motility was scored via thrashing in M9 buffer (22 mM KH_2_PO_4_, 42 mM Na_2_HPO_4_, 85 mM NaCl, 1 mM MgSO_4_). To perform the thrashing assay, L4 animals were first transferred for 5 minutes to an unseeded culture plate to remove bacterial food adhered to their body. The animals were then transferred to 15 μl of M9 buffer on a 15-well multitest slide and allowed to acclimate for 1 minute. Animals were then imaged for 30 seconds using a Moticam X camera (Motic, California, US) mounted to the eyepiece of a Nikon SMZ800 dissection microscope and recorded using a Samsung Galaxy S10 mobile device. Movies were converted from MP4s to AVIs using FFMPEG. The number of body thrashes per 30 seconds was counted for at least 10 animals per genotype.

### Cuticle permeability assay

Cuticle permeability assays using acridine orange were performed as described ^28^. In brief, adult animals were washed off seeded NGM plates and stained with 18.8 μM acridine orange in M9 buffer for 15 minutes. Following three M9 washes, animals were imaged as above. Average fluorescence intensity was measured in Fiji using the polygon tool to outline the uterus in the central uterine plane for 15 animals per condition. Data were analyzed using Student’s t-test in Prism.

### Figures and statistical analysis

Image vignettes were made from images cropped in Adobe Photoshop and assembled in Adobe Illustrator. GraphPad Prism software was utilized to generate graphs and perform appropriate statistical analysis. An analysis of variance (ANOVA) corrected with Tukey Multiple Comparison test or Student’s unpaired t-test determined statistical significance for all figures, as noted in the figure legends. Assumptions for normality and equal variance were made for all analyzed data. A *p* value less than 0.05 was considered significant. All figures report significance testing results and standard deviation error bars. All legends and figures include sample size (*n)* and *p* value.

## End Notes

## Acknowledgments

The authors would like to thank the following individuals for discussions on this work: Lillian Fritz-Laylin, Amy Gladfelter and her lab, Bob Goldstein, Kacy Gordon, Michelle Momany, Brent Shuman, Masayuki Onishi, and Samed Delic. We would also like to thank members of the Maddox labs for comments, particularly Linnea Wethekam for critical reading of this manuscript and Amanda Brown for assistance. Some strains used in this work were provided by the Caenorhabditis Genetics Center (CGC), which is funded by NIH Office of Research Infrastructure Programs (P40 OD010440).

## Author contributions

Conceptualization: JAP

Methodology: JAP, MEW

Investigation: JAP, MEW, SO, BWH, MM

Visualization: JAP

Funding acquisition: JAP, ASM

Supervision: PSM, MM, ASM

Resources: PSM, MM, ASM

Writing – original draft: JAP

Writing – review & editing: JAP, MEW, MM, ASM

## Data and materials availability

All data are available in the main text or the supplementary materials. Information and reagent requests should be directed to the corresponding author(s).

