## Supplementary Materials for "Animal septins contain functional transmembrane domains"

### Supplemental Material

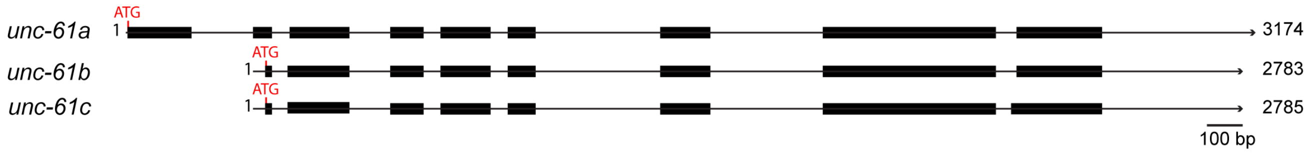


**Fig. S1. *unc-61* locus encodes three isoforms.** A schematic of the *unc-61* locus. Exons are denoted by black boxes; start sites are marked in red; numbers represent base pair.

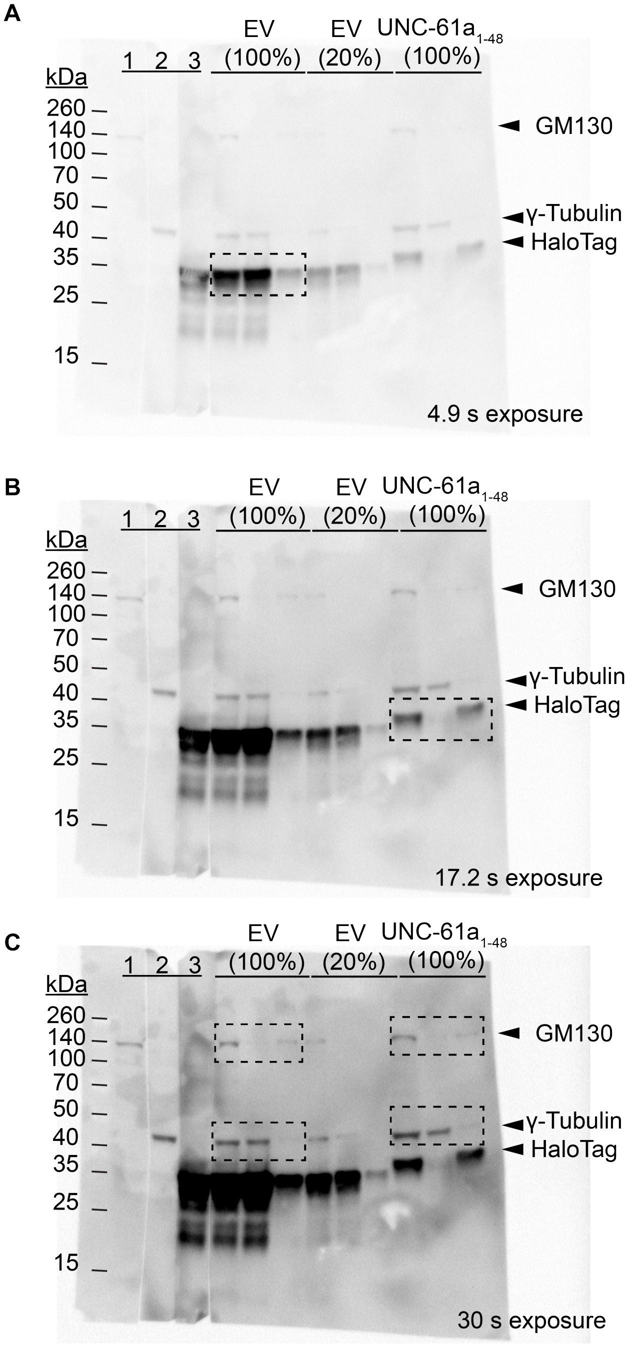


**Fig. S2. Unedited Cell Fractionation Blots. (A)** A 4.9 s exposure of western blot of cytosolic (C) and membrane (M) fractions of HeLa cells transiently expressing either HaloTag empty vector (EV) or UNC-61a $TMD_{1-48}$::HaloTag (UNC-61a $TMD_{1-48}$). Lanes 1-3: whole cell lysate (WCL) blotted singly with antibodies recognizing GM130 (membrane), γ-tubulin (cytosol) and HaloTag. Lanes 4-6 WCL, C, and M fractions are probed with a mixture of all three antibodies (GM130, γ-tubulin, and HaloTag). **(B)** 17.2 s exposure of the same blot as (A). **(C)** 30 s exposure of the same blot as (A). For all blots, dashed boxes indicate the regions used to make Figure 1E.

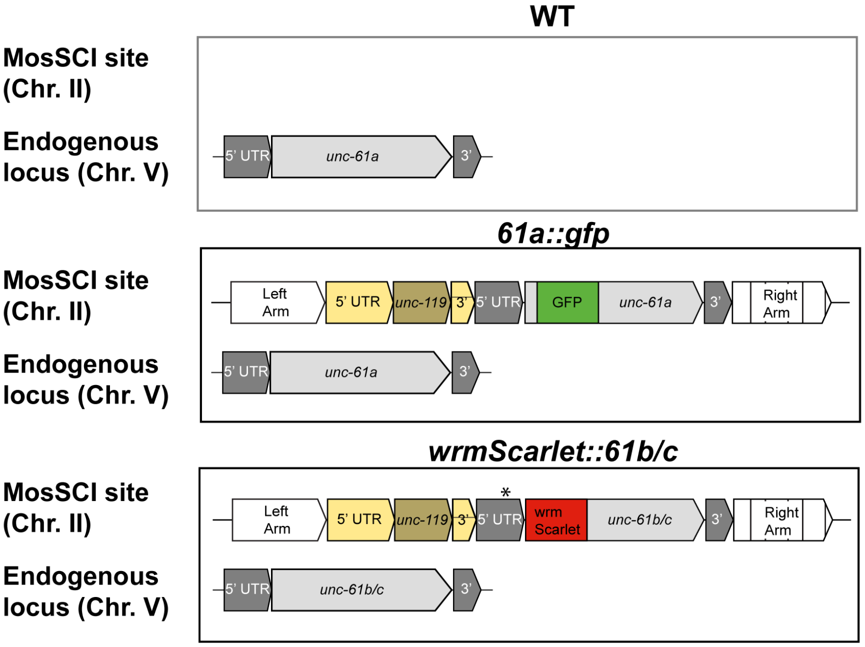


**Fig. S3. Genetic architecture of wild-type, *unc-61a*::GFP, and wrmScarlet::*unc-61b/c*.** Depiction of MosSCI insertion site on chromosome II and *unc-61* endogenous locus (chromosome V). Asterisk (*) denotes location of UNC-61a M1I within the promoter sequence to prevent read-through ofUNC-61a.

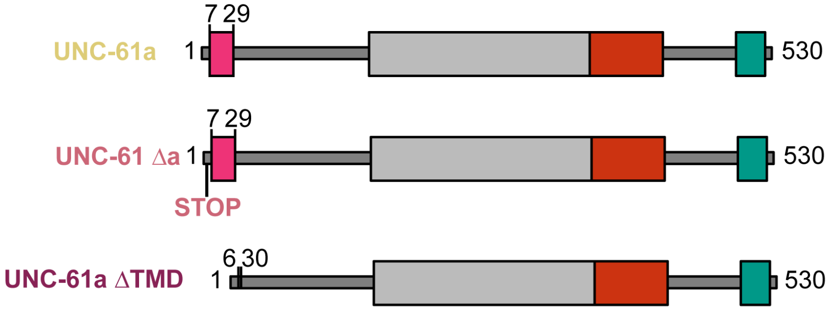


**Fig. S4. CRISPR/Cas9 genome editing scheme for *unc-61a* alleles.** (Top) Schematic of unedited UNC-61a. (Middle) Depiction of *unc-61a* null, where S2* mutation was engineered to introduce a STOP codon right before the TMD. (Bottom) Illustration of UNC-61a ΔTMD where residues 7-29 were excised. Colors as in Figure 1.

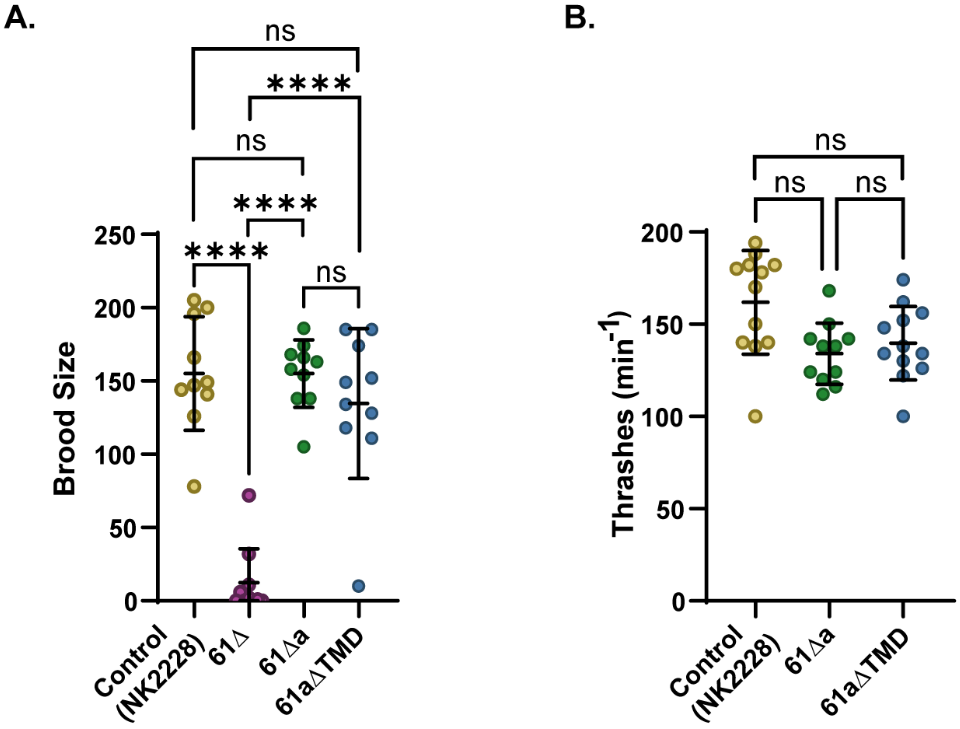


**Fig. S5. Loss of UNC61a or the UNC-61a TMD does not affect animal fertility or motility.** **(A)** Number of embryos produced (brood size) per worm in 48 hours by control, *unc-61null* (*61 Δ*), *unc-61a null (61 Δa), unc-61a ΔTMD (61a Δ TMD)* animals (n = 10). Only *unc-61(null)* animals had significantly reduced fertility (****, p ≤ 0.0001) while the brood size of other animals was not significantly (ns) changed. **(B)** Body thrashes were counted for 30 seconds per worm (n = 10 animals). All data are represented as mean ± SD. Statistical analysis determined by ANOVA and corrected by Tukey Multiple Comparison test.

```
HsSEPT10_i1      MASSEVARHLLFQSHMATKTTCMSSQGSDDEQIKRENIRSLTMSGHVGFESLPDQLVNRS      60
CsSEPT10         ---------------MAMKMTCMSSQVSDDEQRKREKIRSLTMCGHVGFESLPDQLVNRS      45
                                **  * ****** ***** ***:******.****************

HsSEPT10_i1      IQQGFCFNILCVGETGIGKSTLIDTLFNTNFEDYESSHFCPNVKLKAQTYELQESNVQLK      120
CsSEPT10         IQQGFRFNILCVGETGIGKSTLINTLFNTSFEDCESSHFYPHVKLKAQTYELQESNVQLK      105
                 ***** *****************:*****.*** ***** *:******************

HsSEPT10_i1      LTIVNTVGFGDQINKEESYQPIVDYIDAQFEAYLQEELKIKRSLFTYHDSRIHVCLYFIS      180
CsSEPT10         LTVVSTVGFGDQTNKEESYQPIVDYIDAQFEAYLQEELKIERSLFNYLDSRIHVCLYFIS      165
                 **:*.******* ***************************:****.* ************

HsSEPT10_i1      PTGHSLKTLDLLTMKNLDSKVNIIPVIAKADTVSKTELQKFKIKLMSELVSNGVQIYQFP      240
CsSEPT10         PTGHSLKILDLLTMKNLDSKVNIIPVIAQADGISKTELQKFKMALMSELVSNGVQIYQFP      225
                 ******* ********************:** :*********: ****************

HsSEPT10_i1      TDDDTIAKVNAAMNGQLPFAVVGSMDEVKVGNKMVKARQYPWGVVQVENENHCDFVKLRE      300
CsSEPT10         TDDDSVAKINAAMNGQLPFAVVGSMTEVKVGNKMVKARQYPWGVVQVENENHCDFVKLRE      285
                 ****::**:**************** **********************************

HsSEPT10_i1      MLICTNMEDLREQTHTRHYELYRRCKLEEMGFTDVGPENKPVSVQETYEAKRHEFHGERQ      360
CsSEPT10         MLICTNMEDLREQTHSRHYELYRRCKLEEMGFADVGPESQPVSLQQMYDVKRQEFCREVQ      345
                 ***************:****************:*****.:***:*: *:.**:**  * *

HsSEPT10_i1      RKEEEMKQMFVQRVKEKEAILKEAERELQAKFEHLKRLHQEERMKLEEKRRLLEEEIIAF      420
CsSEPT10         RKEEELKRVYVQRVREKEALFKEARRELLAKFEHLKRVHQEETMRLEEKRRSLEEEVIAF      405
                 *****:*:::****:****::***.*** ********:**** *:****** ****:***

HsSEPT10_i1      SKKKATSEIFHSQSFLATGSNLRKDKDRKNSNFL--------------------------      454
CsSEPT10         SKKKVASEVFQNQPFLAVGSNLRKDKDHKKEPSCRFELLCIDVRACEASAGRKDAEKAPI      465
                 ****.:**:*:.* ***.*********:*:.

HsSEPT10_i1      ------------------------------------------------------------      454
CsSEPT10         LCKTEVPDHRKSSQAKAKVIKKLHVCFDFLAVFIYVITYSISGGWPQLVTFMGKCFQVQG      525

HsSEPT10_i1      ------      454
CsSEPT10         QYISQS      531
```

**Fig. S6. Alignment of *Hs*SEPT10_i1 and *Cs*SEPT10.** Sequence alignment of *Hs*SEPT10_i1 and *Cs*SEPT10 using ClustalOmega. The TMD sequence of *Cs*SEPT10 is colored green. The sequence of *Cs*SEPT10$_C$ is underlined. Asterisk (*), colon (:), and period (.) denote level of amino acid conservation.

| STRAIN | GENOTYPE | SOURCE OR REFERENCE |
|---|---|---|
| COP93 | ttTi5606 II; unc119(ed3) III | InVivo Biosciences |
| COP2586 | knuSi921[pNU3401(AMAD02-unc-61p::UNC-61(a)::GFP-C1::unc-61u+, unc-119(+))] II; unc-119(ed3)III. | InVivo Biosciences/This work |
| NK2228 | unc-59(qy50 [unc-59::GFP-C1::;3xflag::AID]) I. | Chen *et al.* |
| MDX130 | unc-59(qy50 [unc-59::GFP-C1::;3xflag::AID]) I; unc-61a(mon24 [unc-61a S2X]) V | This work |
| MDX118 | unc-61(mon22 [unc-61 KO]) V. | This work |
| MDX138 | knuSi921[pNU3401(AMAD02-unc-61p::UNC-61(a)::GFP-C1::unc-61u+, unc-119(+))] II; unc-119(ed3)III ; unc-61(mon22 [unc-61 KO]) V. | This work |
| MDX140 | knuSi921[pNU3401(AMAD02-unc-61p::UNC-61(a)::GFP-C1::unc-61u+, unc-119(+))] II; unc-119(ed3)III; unc-61a(mon24 [unc-61a S2X]) V | This work |
| MDX132 | unc-59(qy50 [unc-59::GFP-c1::;3xflag::AID]) I; unc-61a(mon26 [unc-61a L7_V29del]) V. | This work |
| LP193 | cpIS56[Pmex-5::TAGRFPT:: PLCδ-PH::tbb-2 3'UTR + unc-119 (+)] II; unc-119(ed3) III. | CGC |
| FT1197 | xnls449[lin-26::Lifeact::GFP + unc-119(+)]. | CGC |
| MDX150 | unc-61a(mon26 [unc-61a L7_V29del]) V; xnls449[lin-26::Lifeact::GFP + unc-119(+)]. | This work |
| COP2601 | knuSi934 [pNU3400 (AMAD01-unc-61p:: wrmScarlet::UNC-61(B/C)::unc-61u, unc-119(+))] II; unc-119(ed)III. | InVivo Biosciences/ This work |
| MDX152 | knuSi934 [pNU3400 (AMAD01-unc-61p:: wrmScarlet::UNC-61(B/C):: unc-61u, unc-119(+))] II; unc-119(ed)III; unc-61a (mon26 [unc-61a L7_V29del]) V. | This work |

**Table 1: *C. elegans* strains used in this work**

| PRIMER CODE | PRIMER NAME | OLIGONUCLEOTIDES 5' to 3' | USE |
|---|---|---|---|
| P0285 | NheImsfGFPFOR | CGTCAGATCCGCTAGCATGGTGTCCAAGGGCGAG | pM321, pM324, pM325 |
| P0477 | msfGFPREV | CTTGTACAGCTCATCCATGCCCAGG | pM321, pM325 |
| P0917 | msfGFPSEPT10FOR | GATGAGCTGTACAAGGCCTCCTCCGAGGTGGC | pM321, pM325 |
| P0918 | BamHISEPT10REV | TAGATCCGGTGGATCCTTACAAAAAATTGGAGTTCTTACGGTCCTTGTC | pM321 |
| P1110 | BamHISEPT10iX1tarsierREV | TAGATCCGGTGGATCCTCAACTCTGGCTTATGTATTGACCTTGTAC | pM324, pM325 |
| P1111 | NheImScarletFOR | CGTCAGATCCGCTAGCATGGATAGCACCGAGGCAG | pM300 |
| p1073 | BamHICaaxmScarlet | TAGATCCGGTGGATCCTTAGGAGAGCACACACTTGCAGCTCATGCAGCCGGGGCCACTCTCATCAGGAGGGTTCAGCTTGGAGCCACCGGAGCCGCCG | pM300 |
| JP2415 | CeTMD-pM324-F | TTAGTGAACCGTCAGATCCGCTAGCATGAGTTTCGAAACGATTC | pJAP11 |
| JP2416 | CeTMD-linker-R | TCCGGAGGACCCACCACCTCCAGAGCCACCGCCACCATTCTCATGGGTAACCAC | pJAP11 |
| JP2417 | CeTMD-linker-msfGFP F | GGTGGCGGTGGCTCTGGAGGTGGTGGGTCCTCCGGAGTGTCCAAGGGCGAGGAG | pJAP11 |
| JP2418 | msfGFP-pM324R | TCAGTTATCTAGATCCGGTGGATCCttaCTTGTACAGCTCATCCATGC | pJAP11 |

**Table 2: Primer sequences for construction of mammalian transgenes**

| **PRIMER** | **PURPOSE** | **SEQUENCE** |
|---|---|---|
| CEH13025F | Mutant F | tcgaagatctgcccactagtgagtcg |
| CEH13026R | Mutant R | tgtttgcgtgttctcccattcttcactg |
| CEH13021F | Background F | aggcagaatgtgaacaagactcgagc |
| CEH13023R | Background R | acatacggcaagagcggttcgc |

**Table 3: Primer sequences to confirm MosSCI animals.**

***CRISPR/cas9 Targeting Sequences***

| **GENE** | **TARGETING SEQUENCE** |
|---|---|
| *unc-61* | tttttattcgaactcgatg ; tcttaacttctttgacactt |
| *unc-61a* | tttttattcgaactcgatg |
| *unc-61a* transmembrane domain | tttttattcgaactcgatg ; ccgcttttctcatcgtccga |

***CRISPR/cas9 Repair Sequences***

| **GENE** | **REPAIR SEQUENCE** |
|---|---|
| *unc-61a* null | *ActctgctgaccagacgattttttattcgaactcgatgagAagcacaactgacgaatcgatagataatttATGtAGTTTCGAAACGATTCTCTATACTGCATCTGCCCTTCTT* |
| *unc-61a* transmembrane domain deletion | *actctgctgaccagacgattttttattcgaactcgatgagTagcacaactgacgaatcgatagataatttATGAGTTTCGAAACGATTCGACGATCGAAACAAAGCAGCATAAACCTGCAGAC* |

***Primer sequences***

| **PRIMER** | **SEQUENCE** |
|---|---|
| UNC-61 null 5' PCR Primer | gacagttgttggttatatttgc |
| UNC-61 null 3' PCR Primer | tgggtgtagtgtgtatttgaaa |
| UNC-61a deletion 5' PCR Primer | ccgcttttctcatcgtccga |
| UNC-61a deletion 3' PCR Primer | gctgcaattgttggcttcttgc |
| UNC-61a transmembrane domain deletion 5' PCR Primer | ccgcttttctcatcgtccga |
| UNC-61a transmembrane domain deletion 3' PCR Primer | gctgcaattgttggcttcttgc |

**Table 4: Reagents to construct and confirm *C. elegans* CRISPR/Cas9 edited strains**
